# Pupil size as a robust marker of attentional bias toward nicotine-related stimuli in smokers

**DOI:** 10.1101/2022.05.08.490888

**Authors:** Elvio Blini, Marco Zorzi

## Abstract

Spatial attention can be magnetically attracted by behaviorally salient stimuli. This phenomenon occasionally conflicts with behavioral goals, leading to maladaptive consequences, as in the case of addiction, in which attentional biases have been described and linked with clinically meaningful variables, such as craving level or dependence intensity. Here, we sought to probe the markers of attentional priority in smokers through eye-tracking measures, by leveraging the established link between eye movements and spatial attention. We were particularly interested in potential markers related to pupil size, because pupil diameter reflects a range of autonomic, affective, and cognitive/attentional reactions to behaviorally significant stimuli and is a robust marker of appetitive and aversive learning. We found that changes in pupil size to nicotine-related visual stimuli could reliably predict, in crossvalidated logistic regression, the smoking status of young smokers (showing pupil constriction) better than more traditional proxy measures. The possibility that pupil constriction may reflect a bias toward central vision, e.g. attentional capture, is discussed in terms of sensory tuning with respect to nicotine-related stimuli. Pupil size was more sensitive at lower nicotine dependence levels, and at increased abstinence time (though these two variables were collinear). We conclude that pupillometry can provide a robust marker for attentional priority computation and useful indications regarding motivational states and individual attitudes toward conditioned stimuli.

## 1. Introduction

Intrinsically rewarding stimuli attract spatial attention (Anderson et al., 2011; Awh et al., 2012; Chelazzi et al., 2013; Hickey et al., 2010; Sprague & Serences, 2013). Priority maps in the brain represent the physical, low-level properties of stimuli but, the higher the hierarchy level, the more maps start to encode behavioral relevance beyond those features (Bisley & Mirpour, 2019; Fecteau & Munoz, 2006). Traces of behavioral salience can be found in the brain as early as the superior colliculus (White et al., 2017a; White et al., 2017b) and the primary visual cortex (Li, 2002), although still confined to the physical features that they encode (Bisley & Mirpour, 2019). The concept of priority maps helped moving beyond classic dichotomies (i.e., bottom-up vs. top-down attention, Awh et al., 2012; see Anderson, 2020). Value-driven attentional capture (Anderson et al., 2011, 2014; Bourgeois et al., 2016), on the one hand, and statistical learning or selection history (Duncan & Theeuwes, 2020; Failing & Theeuwes, 2018; Jiang et al., 2015), on the other hand, appear to escape these labels, all while representing powerful determinants of attentional biases. This is arguably efficient and evolutionarily convenient, in that both attention and reward serve to filter the most relevant sensory information as to optimize cognitive adaptation (Driver, 2001; Manohar et al., 2015; Maunsell, 2004). As such, they are strictly intertwined processes, to the point that they can easily be confounded (Anderson et al., 2011; Maunsell, 2004; Raymond & O’Brien, 2009). Yet, studying them, and their interaction, is not only relevant for understanding human cognition, but also provides a framework to study instances in which value conflicts with behavioral goals, leading to maladaptive consequences, as in addiction.

Addiction disorders are multifaceted conditions. Among their many features, dysfunctions to the processing of rewards have been described, and consist of an increased sensitivity to selected reinforcers at the expense of alternative ones (Goldstein & Volkow, 2002; Noël et al., 2013). These stimuli gain high attentional priority in everyday life situations, creating a vicious loop believed to help maintaining addiction (Volkow et al., 2010), and to the point of hampering inhibitory and monitoring processes (Blini et al., 2020; Cox et al., 2006; Field et al., 2007; Field & Cox, 2008; Goldstein & Volkow, 2011; Noël et al., 2013). Not surprisingly, interventions aimed at modifying attentional biases have been proposed. Their goal is to divert attention away from disorder-relevant stimuli, and this is chiefly achieved by leveraging on statistical learning procedures. However, the effectiveness of these interventions remains to date, inconclusive (Cox et al., 2014; Mogoaşe et al., 2014). While the causal role of attentional bias in addiction is still debated, however, attentional bias itself remains a valid marker of the underlying, fluctuating motivational state (Christiansen et al., 2015). Even if considered a byproduct, thus, it can still represent a useful proxy for clinically meaningful information such as current craving level (Field et al., 2014), current concerns about addiction, or the broad affective reaction to these meaningful stimuli. As such, it is important to deepen our knowledge and characterization of attentional biomarkers.

An established paradigm to study attentional biases is the Dot-Probe Task (DPT; Ehrman et al., 2002; Waters et al., 2003). In the case of smokers, two pictures are shown on either side of a computer screen, one depicting a Nicotine-Related Stimulus (NRS), one a physically matched neutral object. After a variable delay, and along the tradition of Posner-like cueing tasks (Posner, 1980), one to-be-discriminated target is presented on either side. Active smokers present a response times advantage whenever the target is presented on the same side of the NRS, showing that spatial attention is magnetically attracted by the cue (Ehrman et al., 2002). This effect appears to go beyond the increased familiarity of smokers with NRS (Chanon et al., 2010), and it is highly dependent on Stimulus-Onset Asynchronies (Della Libera et al., 2019). Crucially, it is stronger for direct measures of attention (Field et al., 2009). Electroencephalographic studies, for example, have shown that masked NRS elicit a larger N2pc component (that is, an index of lateralized spatial attention) than masked control stimuli (Harris et al., 2018), suggesting that the former, at least in conditions of reduced perceptibility, capture visual attention more strongly very early (i.e., around 250 ms). Another important direct proxy of attentional bias is eye movements, because eye movements and spatial attention – whilst dissociable (e.g., Weaver et al., 2017) – are profoundly linked (Blini, Pitteri, & Zorzi, 2019; Casarotti et al., 2012; Rizzolatti et al., 1987). When stimuli are given in the visual modality, the oculomotor system codes and is biased by their value (Camara et al., 2013; Manohar & Husain, 2015; Muhammed et al., 2020; Takikawa et al., 2002; Theeuwes & Belopolsky, 2012). This is also seen, in smokers administered with the DPT, as longer time spent fixating NRS and/or first fixations more often directed toward these stimuli (Field et al., 2004; Mogg et al., 2003, 2005).

Pupil size is a more special measure among those that can be inferred by recording the eyes. Pupil diameter changes first and foremost following changes in the amount of light entering the eyes (see, e.g., Binda & Murray, 2015). However, it has been associated with phasic activation of the locus coeruleus (Aston-Jones & Cohen, 2005), a major noradrenergic hub involved in the integration of the attentional systems in the brain, as well as in balancing bottom-up and top-down aspects of perception (Reynaud et al., 2021). Pupil diameter reflects, beyond that, the activation of the autonomic system, and as such it ideally probes: affective and emotional responses to visual stimuli (Bogdanova et al., 2022; Dureux et al., 2021); the amount of cognitive effort deployed for a task (e.g., working memory load, Lisi et al., 2015; Mathôt, 2018); the subjective value of a given stimulus (Muhammed et al., 2016; Pietrock et al., 2019). Accordingly, a recent meta-analysis concluded that pupil dilation is a robust marker of Pavlovian-like conditioning (Finke et al., 2021). Despite these premises, however, studies attempting to characterize attentional biases in addiction by means of pupillometry are still scarce (but see Chae et al., 2008). The current study aims to fill this gap, by assessing: i) whether pupil can, in a fast event-related design, reliably assess attentional biases toward NRS; ii) how this measure compares with more common ones (e.g., as obtained from a DPT task); iii) whether it can be used for a reliable classification of smokers; iv) whether it correlates with either smoking intensity or craving urges.

## 2. Methods

All materials, raw data, and analysis scripts for this study are available at the following link (referred to as Supplementary materials): https://osf.io/6r5ch/

### 2.1 Participants

In this work, we planned to acquire multiple measures of attentional and autonomic engagement with Nicotine-Related Stimuli (NRS) in smokers vs. controls. Instead of calibrating the recruitment plan on each individual measure, we reasoned that a multivariate approach – attempting to classify the group on the basis of these measures – would be a more appropriate and powerful approach. Thus, we conducted an a priori power analysis for a one-sided binomial test (i.e., testing that classification performance is superior to chance level). We assumed the minimum effect size of interest (and practical relevance) to be a classification accuracy of 70%. With a type 1 error rate set to alpha= 0.05, 80% statistical power for this design is reached at N= 40 (**Fig. 1**). Power analysis was conducted through simulations, and the relative scripts are available in the Supplementary Materials. Note that, at N= 40, a classification accuracy of 65% would be significantly superior to chance (p= 0.04) according to this test; however, this is to be considered the underlying, “true” effect size, which does not invariantly lead to the same classification accuracy due to random fluctuations, hence the concept of statistical power of a test. Therefore, we planned to enroll a minimum of 20 smokers and 20 matched controls. The assumption of our elective multivariate classifier was that all participants must have all the pre-identified behavioral predictors, without missing data. Based on our set of data quality (explained below), this led to the recruitment of 51 participants in total.

**Fig. 1.**
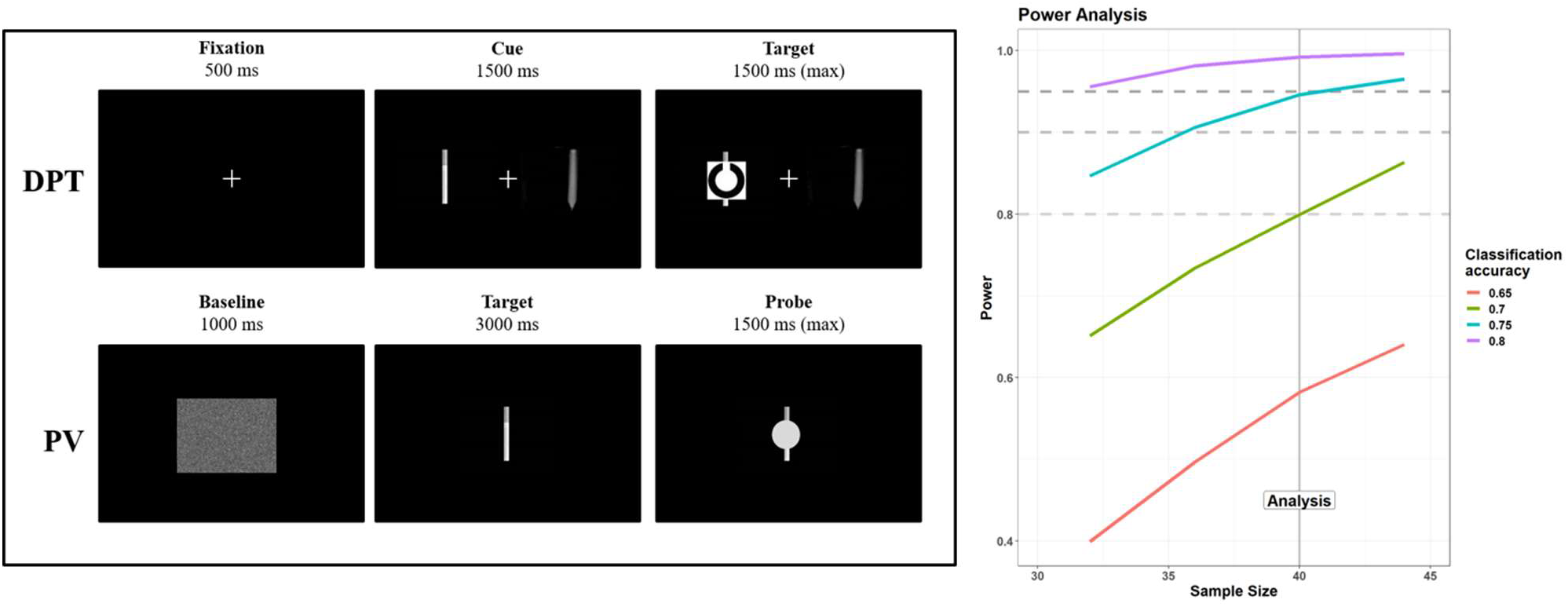
Left panel: structure and time course of the two experimental tasks, Dot-Probe task (DPT, upper row) and Passive Viewing (PV, bottom row). DPT) Each trial started with a fixation cross, presented for 500 ms. Then, two pictorial stimuli were presented on each side of the screen, one depicting a Nicotine-related Stimulus (NRS) and one a matched, control image. The stimuli remained on screen for a relatively long SOA, 1500 ms, during which eye fixations were recorded. Then, one to-be-discriminated target appeared on either side, and response times were collected. PV) Each trial started with a baseline visual stimulus obtained by randomly shuffling pixels in the upcoming images. Then, either NRS or control images were presented, foveally, for 3 seconds; participants were instructed to passively look at the images. In a minority of trials (30%) one probe appeared on screen after the 3 seconds window: the probe had to be detected as fast as possible. Note that stimuli depicted in the image are only examples, both tasks involved pictures of people smoking (see main text). An overview of all stimuli is available in the supplementary materials (https://osf.io/6r5ch/). Right panel: nominal power for our design depicted as a function of sample size and varying degrees of a priori effect sizes (i.e., classification accuracy to a binomial test). The gray lines depict three conventional statistical power thresholds: 80%, 90%, 95% power.

The sample was composed of 28 smokers (age: 24.04 ± 2.69 years; 18 were women, 4 were left handed) and 23 non-smokers (age: 23.13 ± 2.75 years; 16 were women, 1 was left handed). All participants were recruited from students of the University of Padova (Italy). Inclusion criteria were normal (or corrected-to-normal) vision and no history of neurological or psychiatric diseases. Occasional (i.e., non-daily) smokers were excluded and all smokers in the final sample smoked a self-reported mean of 8.1 cigarettes a day and for an average duration of 5 years. Most of them were light smokers, however, as seen by low average scores (1.96 ± 2.1) at the Fagerström test for Nicotine Dependence (FTND; Heatherton et al., 1991). Because nicotine can considerably reduce pupil size (Wardhani et al., 2020), and because we sought to evaluate the tasks in conditions of relative craving, they were all asked to refrain from smoking for at least 90 minutes prior to the experiment (mean: 7.5 hours; indeed, half of them last smoked the evening before the experiment). In addition to the FTND, smokers were asked to fill in the Severity of Dependence Scale (SDS; Gossop et al., 1992) and the Questionnaire of Smoking Urges (QSU-brief; Cox et al., 2001), to assess self-reported craving. The study was approved by the Ethics Committee of the University of Padova (protocol n° 3568). It was carried out during a period of restrictions due to the covid-19 epidemic and as such all the necessary sanitary precautions were taken.

### 2.2 Materials and Methods

Participants were tested in a dimly lit, quiet room, their head comfortably restrained by a chinrest. They faced a remote infrared-based eye-tracker (TOBII™ Spectrum), with an embedded 24 inches monitor, at a distance of approximately 57 cm. The session started with a 9-points calibration of the eye-tracker, which was then set to monitor participants’ gaze continuously at a 600 Hz sampling rate. The open-source software OpenSesame (Mathôt et al., 2012) was used to display experimental stimuli on the screen and record the subjects’ responses. Participants provided responses by means of keyboard presses (on a standard QWERTY keyboard) using the index and middle fingers of their dominant hand. All participants performed two tasks, in a fixed order: Dot-Probe Task (DPT) and Passive Viewing (PV) task (**Fig. 1**).

#### DPT

The first task was meant to measure spontaneous (spatial) attentional biases by nicotine-related stimuli (NRS). Each trial started with a fixation cross appearing at the center of the screen for 500 ms. Then, two images (6° x 4.5°) were always presented concurrently on the left and right sides of the screen, about 2° away from fixation horizontally. One image depicted a Nicotine-Related Stimulus (NRS; e.g., a person smoking), and one a perceptually-matched control stimulus (e.g., a person with otherwise similar features, see the paragraph below). The side of NRS appearance (i.e., left, right) was equiprobable. We choose a relatively long on-screen duration for these images (i.e., SOA, 1.5s). Previous studies have highlighted how attentional biases can unfold at different temporal windows, e.g. differently at 200 ms than 800 ms (Della Libera et al., 2019). Using only one, extra-long SOA is guaranteed to miss this fine-grained dissociation. However, our choice was motivated based on pilot testing and ultimately our stress on overt attentional measures (i.e., eye movements) over response times, which have been found to be more sensitive to craving (Field et al., 2009) although requiring an extended time window (Field et al., 2004; Mogg et al., 2003). After this long SOA, the target appeared either superimposed on top of the NRS or the control image (balanced). The target was a circle (1.5° diameter) with a gap either in its upper or lower part, and participants had to indicate the position of the gap through the corresponding arrow keys. The target remained on screen until the participants’ response, up to a maximum of 1.5s. Participants were first administered 12 practice trials, later discarded from analyses, followed by 240 experimental trials. A break was foreseen halfway through the task in order to allow participants to rest.

#### PV

The second task was a passive viewing task of NRS or control images presented centrally on the screen (4.8° x 3.6°), and the main objective was to acquire indices of autonomic activation through patterns of pupil dilation and constriction. Each trial started with 1s foveal presentation of a scrambled version of the target image; then, each image was presented on screen for 3s. Pupil diameter was measured continuously in both these phases. In a minority of trials (30%), a visual cue (a gray dot, 1° diameter) appeared soon after the presentation of the target image (hence, after 3 s of image presentation, not jittered). The aim was to encourage sustained attention to the images, in absence of an otherwise active behavioral task. The cue remained on screen until the participants’ detection (response given with the spacebar) up to a maximum of 1.5 s. During the inter-trial interval, we also presented scrambled images on screen, as to keep all sources of luminance constant. Participants were first administered 8 practice trials, later discarded from analyses, followed by 200 experimental trials. A break was foreseen halfway through the task in order to allow participants to rest.

#### Images selection

Images were drawn from the SmoCuDa database (Manoliu et al., 2020), a validated and well-described collection of smoking-related images. We selected 10 images, all depicting people smoking (i.e., social stimuli). Each stimulus was paired, to the best of our possibilities, to a control image, matched for all the relevant features but the smoke-related content. Images were all originally 800 x 600 pixels large, before being rescaled to fit the needs of each task. All stimuli were transformed into grayscale images, and were processed in order to have the same mean luminance. The 600×800 matrices of values (between 0 and 1) were first z-transformed; then, each picture was transformed again to a new, common distribution of values having a mean intensity of 0.4 and a standard deviation of 0.15. For each image, a scrambled version was obtained offline by randomly shuffling the indices of the final grayscale matrices. The resulting images had the same mean luminance, but were hardly recognizable; note that, however, we did not in fact surveyed our participants for this or for the possibility of learning effects. Images can be assessed in the supplementary materials (https://osf.io/6r5ch/).

### 2.3 Data processing

#### DPT

We discarded practice trials and anticipations (responses faster than 100 ms, <0.005%). All participants performed well in this task (minimum accuracy: 80.4%), thus, there were no exclusions related to this measure of behavioral performance. Furthermore, because accuracy was on average very high (M= 98.4%, SD= 2.95%), with little variability, this measure was not further considered in the analyses. We analyzed, instead, the mean Response Times (RTs) for correct responses. We computed RTs for each Condition (NRS, control) and then subtracted the former from the latter so that positive values index enhanced attentional engagement with NRS (i.e., faster responses when probes appear over NRS).

We then moved to assess eye movements. We only retained X and Y positions of the gaze when data were available for both eyes, and took their average value. As a cautionary measure, we discarded the trials in which more than 40% of the data were missing (this includes blinks, artifacts, lost connection with the eye-tracker, etc.); furthermore, we established, prior to testing, to discard from the sample participants remaining with fewer than 50% of valid trials. This led to the exclusion of 6 participants from the main analysis (note: other participants, for a total of 11, were discarded for the same reason applied to the PV task). Missing trials for the remaining participants were very few (for smokers: 2.5% ± 5.2%; for controls: 5.2% ± 8.2%). The gaps in the remaining recordings were then linearly interpolated for both the X and Y axes. We focused the analyses on the 1500 ms window in which the cue was presented, prior to the target probe. We reconstructed, in this window, the pattern of eye fixations through an automated, velocity-based algorithm (von der Malsburg, 2015), and only considered fixation events lasting at least 40 ms. As a first measure, we computed, for each participant, the proportion of first fixations falling within the NRS region vs the control region. This variable was meant to index a very early, automatic capture of attention and eye movements by NRS. Then, we computed dwell time, for each trial, as the cumulative duration of fixations in each image Condition. This variable is also in line with previous studies, and was meant to index a more or less thorough visual scanning of the different image classes. Both these variables were transformed into bias scores by subtraction so that negative values reflect a bias towards control images, and positive towards NRS.

#### PV

We first discarded practice trials. Missed responses to the infrequent probe were very rare (0.002%), showing some engagement with the presented images. As a first measure, we computed the RTs to the probe for NRS vs. control images, and then subtracted the former from the latter so that positive values index enhanced attentional engagement with NRS.

We then moved to assess the time course of changes in pupil size, mostly capitalizing on preprocessing steps adopted in previous studies (Dureux et al., 2021). First, we took the average pupil diameter of the left and right eyes, but only when both signals were properly recorded (according to the eye-tracker’s built-in criteria). We focused our analyses on the baseline period (1s, scrambled images) and the 3s window in which images were presented. Implausible values for pupil diameter (<2 mm and > 7mm), although infrequent (<0.0002%), were omitted. For missing samples within each trial, we followed an interpolation strategy akin to that used for the DPT: we discarded trials in which more than 40% of the data were missing (this includes blinks, artifacts, lost connection with the eye-tracker, etc.), and linearly interpolated gaps in the remaining ones. Participants with fewer than 50% of valid trials (N= 11) were discarded from the main analyses. Missing trials for the remaining participants were few (for smokers: 8.4% ± 13.6%; for controls: 10% ± 13%). We applied a low-pass filter to the raw traces and ultimately down-sampled the data to 25 ms epochs by taking the median pupil diameter for each time bin. The average pupil diameter was similar in the two groups (4.7mm for smokers vs 4.56mm for controls, t_(37.66)_= 0.462, *p*= 0.647). Note that the acute effects of nicotine intake are known to globally constrict the pupils (Wardhani et al., 2020), but we asked all participants to refrain from smoking prior to the experiment. At any rate, in order to better cope with individual (and group) differences we z-tranformed pupil diameter values separately for each participant (Dureux et al., 2021). This way, a value of 0 represents the subject-specific mean pupil diameter and, regardless of baseline values, scores represent dilation (positive values) or constriction (negative values) expressed as a fraction of the overall participant’s variability. Finally, all series were realigned to the beginning of the target image phase (either NRS or control) via subtraction of the first sample in this epoch. We assessed the traces in full when analyses focused on the PV task alone. For the main analyses, however, exploiting a multivariate classifier, we decided to avoid the extremely high dimensionality of these data, which yields overfitting, and rather focused on a time window identified by a cluster-based permutation test. Cluster-based permutation test can handle very elegantly the problem of multiple comparisons in autocorrelated data, although this may come at the price of a more imprecise estimation of the temporal features of the reported effects (e.g., latency, see Sassenhagen & Draschkow, 2019); however, since pupil size is a much slower physiological signal with respect to, for example, the electric or magnetic signals captured by EEG or MEG, we did not anticipate this parameter to be critical. Traces from NRS vs control images were subtracted so that positive values indexed enhanced pupil dilation to NRS, and negative values a relative constriction (see **Fig. 2**); the values used as predictors were therefore the mean, cluster-wide differences in these curves for each participant.

**Fig. 2.**
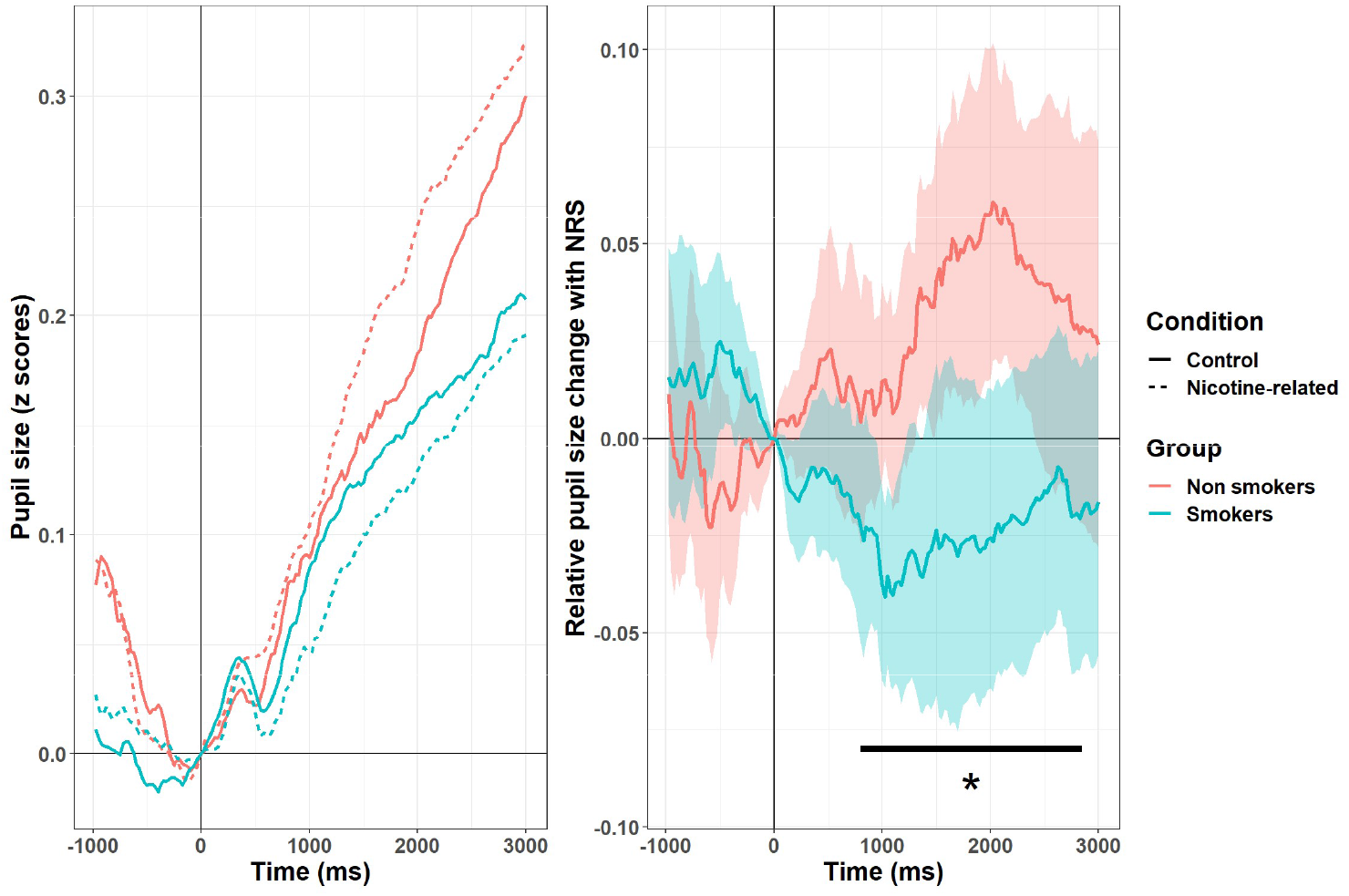
Time course of changes in pupil size during the baseline and target presentation windows (left panel); in both panels positive values index dilation, and negative values constriction of the pupils. Pupils dilated steeply over time, likely due to the expectation of the appearance of a visual probe, and the relative motor preparation. Differences between NRS and perceptually matched images, however, were not the same in the two groups (right panel: shadowed areas represent 95% confidence interval for the data). Cluster-based permutations highlighted, in a data-driven fashion, a rather extended window (800-2850 ms, highlighted in the plot with a black horizontal line) in which smokers presented, on average, a pattern of pupil constriction to NRS when compared to non-smokers.

### 2.4 Analyses

Analyses were performed with R 4.1.2 (The R Core Team, 2018). Each measure of interest was first analyzed individually by means of two-sample t-tests with Welch’s correction for unequal variances (two-tailed). The main interest of this work, however, was to probe the generalizability of these measures and their ability to predict smoking status of out-of-the-box, new individuals. This was to probe the global capability of eye-tracking and autonomic measures to generalize and possibly represent a viable biomarker of smoking behavior. We started with computing general (logistic) linear models with a Best Subset Regression (BSR) approach. All possible combinations of the predictors were tested in glms, and the overall best model was selected only based on the lowest Bayesian and Akaike Information Criteria, which mitigates the problem of multiple comparisons. This approach was used for the initial features selection step, and the best features were then probed in follow-up crossvalidated logistic regressions. We adopted a Leave-One-(Subject)-Out (LOO) crossvalidation setup, where just one participant was circularly included in the test set. Then, we computed sensitivity, specificity, and area under the curve of the classifier, together with an overall classification accuracy measure, which was submitted to a one-tailed binomial test.

Finally, we performed correlations between the variables of interest, obtained from the two tasks, the individual measures related to the intensity of smoking (e.g., number of cigarettes per day) or subjective craving (e.g., QSU-brief, abstinence duration), and the predictions of the classifier for each participant (i.e., the estimated probability to belong to one group or another). For this last part, we only report a selection of the results, for brevity, although the results can be assessed in full in the supplementary materials. These analyses should be taken with caution in light of the small sample size considered.

## 3. Results

### 3.1 Individual results

Descriptive results are reported in **Table 1**.

**Table 1.** Mean (standard deviation) descriptive values for the five variables of interest, separately for group and image condition (NRS: nicotine-related stimuli).

| Task | Variable | Non-smokers |  | Smokers |  |
| --- | --- | --- | --- | --- | --- |
|  |  | Control | NRS | Control | NRS |
| DPT | RTs (ms) | 540(69) | 537(71) | 547(88) | 538(87) |
|  | Dwell time (ms) | 463(118) | 468(116) | 502(85) | 513(84) |
|  | First Fixation (%) | 51(3.2) | 49(3.2) | 50(2.9) | 50(2.9) |
| PV | RTs (ms) | 416(68) | 420(70) | 448(106) | 448(105) |
|  | Pupil size (z scores) | 0.175(0.148) | 0.211(0.137) | 0.140(0.104) | 0.116(0.083) |

#### DPT

Of the three variables considered for this task, none differed between groups. RTs: *t*_(36.59)_= 1.51,*p*= 0.139, two-tailed. Dwell time: *t*_(28.89)_= 0.485, *p*= 0.631, two-tailed. Proportion of first fixations: *t*_(37.88)_= 0.169, *p*= 0.867, two-tailed.

#### PV

RTs did not differ between the two groups when the probe appeared over NRS vs. control images (t_(37.82)_= 0.55, *p=* 0.58, two-tailed). The assessment of pupil size, however, revealed significant group differences. The changes in pupil diameter were overall dominated by a sustained, steep pattern of pupil dilation, but there was an interaction between Group and Condition as well (**Fig. 2**). Cluster-based permutation test revealed a large temporal cluster (800-2850 ms) in which pupil size to NRS vs control images significantly differed between groups (p<0.001, 5000 permutations). We thus computed the mean, cluster-wide difference between pupil diameter to NRS vs control images for each participant between 800 and 2850 ms. In this window, smokers presented a pattern of overall constriction of the pupil (M= −0.025, SD= 0.075) which, when compared to the overall pattern of pupil dilation observed in non-smokers (M= 0.036, SD= 0.067) resulted significantly different (t_(37.58)_= 2.7, *p*= 0.01, two-tailed). This variable was then used for classification. We also repeated, for each group, the cluster-based permutation test, looking for time points with a significant dilation or constriction (instead of a difference between the other group). There was a temporal cluster (925-1175 ms), in smokers, in which the pupil constricted significantly for NRS images (t_(19)_= −2.83, *p*= 0.01). A significant dilation, instead, was observed for non-smokers later on (1475-2375 ms, t_(19)_= 2.75, *p*= 0. 013). There were no other significant deviations from baseline.

### 3.3 Multivariate classification

BSR modeling of smoking status selected the logistic model with only one predictor, pupil size, as the best compromise between complexity and fit to the data. The BIC of this model was 55.57 (AIC= 52.2), far better than the null model (BIC= 59.14, AIC= 57.45), but only marginally better than the model also including RTs to the DPT task (BIC= 57.5, AIC= 52.43). Because the models were close, we kept both these two features for follow-up, crossvalidated logistic regressions.

The best model included both pupil size to NRS images and RTs: crossvalidation accuracy reached 75%, with very good sensitivity and specificity (both 75%), and AUC of 0.745. The performance was significantly superior to chance according to a one-tailed binomial test (p= 0.001, CI_95_%[58.8% – 87.3%]). The second best model only included pupil size as predictor of smoking status. The performance of the classifier in this case was good, with an accuracy of 65% (both sensitivity and specificity were 65%, AUC was 0.723). This performance was also significantly superior to chance according to a one-tailed binomial test (p= 0.04, CI_95_%[48.3% – 79.4%]). When compared against a chance level of 65%, the model including both predictors was not significantly better (p= 0.12). For comparison, the model only including RTs performed very poorly, with an accuracy in crossvalidation of 52.5% (Sensitivity: 55%; Specificity: 50%; AUC: 0.535).

Overall, these results suggest very good classification capability for measures of attentional bias and in particular the autonomic response conveyed by the pupil, which appears necessary and sufficient for classification of smoking status.

### 3.3 Explorative analyses and correlations

Two measures from the DPT task were correlated with pupil size in the subsequent PV task. This suggests that the observed pattern of pupil constriction in smokers may be partially related to behavioral measures that have been more classically associated with attentional bias. First, there was a negative correlation with dwell time (r= −0.42, CI_95_%[−0.65, −0.12], t_(38)_= 2.83, *p*= 0.0073), which was of similar magnitude in the two groups. A more thorough visual exploration of NRS images was correlated with a more vigorous *constriction* of the pupils in the PV task. Second, there was a positive correlation between pupil size to NRS and the proportion of first fixations toward NRS stimuli in the DPT task (r= 0.29, CI_95_%[-0.02, 0.55], t_(38)_= 1.89, *p*= 0.066). When assessed within each group, the correlation was significant for smokers only (smokers: r= 0.54, CI_95_%[0.12, 0.79], t_(18)_= 2.69, p= 0.015; non-smokers: r= 0.07, p= 0.76, n.s.), though the difference between correlations did not reach statistical significance (Fisher’s z= 1.533, p= 0.125). Smokers presenting a more vigorous pupil constriction in the PV were those with a more pronounced tendency to direct the initial fixation *away* from NRS. Overall, dwell time and the direction of the initial fixation were positively correlated in non-smokers (r= 0.46, CI_95_%[0.02, 0.75], t_(18)_= 2.186, *p*= 0.042), but not in smokers (r= −0.02, *p*= 0.95, n.s.).

We then moved our focus to the group of smokers. We first observed that all the three questionnaires were correlated among them, though not with the behavioral measures drawn from the tasks. We assessed the classifier’s predictions for each smoker – i.e., the probability to belong to the smokers’ group vs the non-smokers group (values <0.5, i.e. classification errors). Surprisingly, the classifier performed best (and the pupil constricted more vigorously) for participants who smoked less (r= −0.48, CI_95_%[−0.76, −0.05], t_(18)_= 2.33, *p*= 0.03). We also note that there was a correlation between the classifier’s performance and abstinence time, i.e. hours since the last cigarette. The longer the abstinence, the better the classification (r= 0.55, CI_95_%[0.15, 0.8], t_(18)_= 2.82, *p*= 0.01). In other words, false negatives were more common in individuals who smoked more recently (**Fig. 3**).

**Fig. 3.**
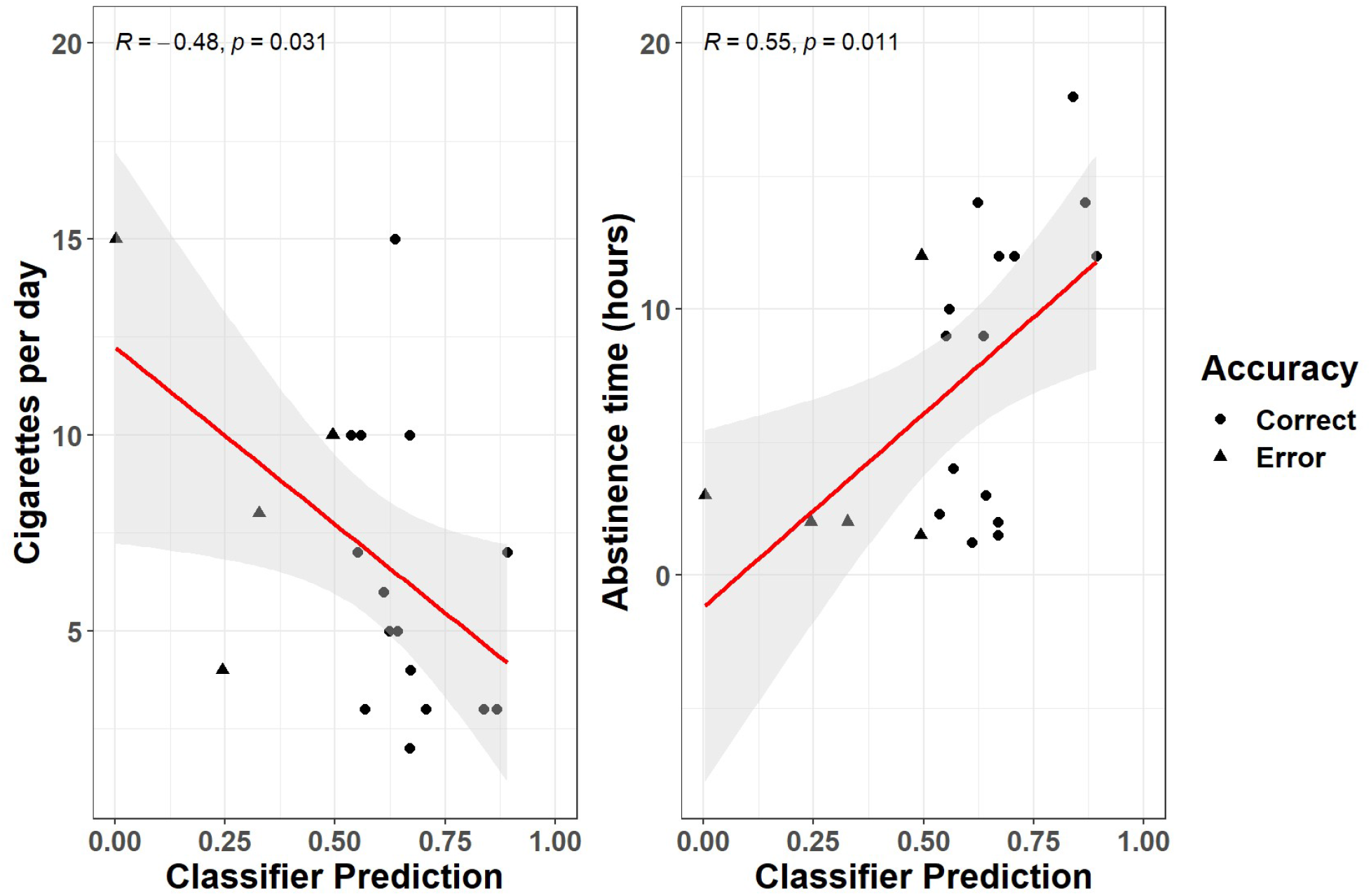
The relation between the classifier’s performance (data points are either circles, i.e. the classification was correct, or triangles, i.e. the classification was not correct), and measures of smoking intensity (self-reported cigarettes per day) and abstinence time (hours). Data points only depict smokers, thus errors are all false negatives. The classifier’s predictions were mostly driven by pupil size, although there was a minor role for RTs in the DPT task as well. The classifier worked best, and the pupil constricted more vigorously, with light smokers and with participants who refrained from smoking for longer periods prior to the experiment (which are partly the same subgroup of participants).

## 4. Discussion

Here, we sought to probe the markers of attentional priority in smokers through eye-tracking. We did not find that behavioral and eye fixation measures were sensitive enough to index attentional biases. However, we found differences in the time course of pupil size changes to NRS between smokers and non-smokers, which proved to be a reliable predictor of smoking status. We used a passive viewing task in which participants were asked to fixate NRS or neutral images for 3 seconds. However, to ensure minimal engagement with all images, rare probes were presented in a minority of trials, after the presentation of images, and participants had to quickly detect them. As a result, the main pattern observed at the level of pupil diameter was a very large, steep, and sustained dilation. This can be readily interpreted in terms of mounting temporal expectations about the appearance of the probe, and the inherent preparation for the motor response (Akdoğan et al., 2016; Wang et al., 2016). We additionally found an interaction, starting from about 800 ms and sustained up to 2850 ms, in which smokers’ pupils tended to *constrict* more than non-smokers ones with NRS. This is interesting, albeit not entirely expected on the basis of evidence pointing to a robust role of pupil dilation in both appetitive and aversive conditioning (Finke et al., 2021). The only previous study that assessed pupil size in response to NRS (Chae et al., 2008) reported pupil dilation in a sample of 7 smokers (against 12 controls). However, their task was a pure passive viewing task in which images were presented on screen for a very long duration (30s). Because our time window was more restricted, we cannot interpolate the results beyond 3s, and the possibility of dilation occurring at later stages remains valid; on the other hand, the fast event-related design of our study may have enabled an unprecedented precision in the earliest perceptual phase. While this is not the only explanation, we speculate that results may be ascribed to an early attentional orienting toward NRS in smokers. The main reason why pupils constrict is light and the Pupillary Light Reflex (PLR), which starts around 200-250ms and can be protracted up to 1-2 seconds (Mathôt, 2020). Even when luminance is strictly controlled for, PLR may still occur, contingently on a change in the visual scene, depending on several factors like awareness, eye-movements, or visual attention (Mathôt & van der Stigchel, 2015). We speculate that NRS stimuli could lead, in smokers, to a pattern of eye-movements or visual attention that is more likely to magnify the PLR in certain images. The resulting pupil responses can indeed be regarded as predictive sensory tuning processes: constriction signals a bias in favor of central vision, since smaller pupils bring about better visual acuity (Mathôt, 2020). A second alternative concerns the specific task demands of our paradigm. The main task here was to react quickly to the infrequent probe, and the pupil was chiefly reflecting that with a strong dilation; relative constriction in this setting may suggest that attentional resources were diverted from the main goal. In other words, the constriction we describe here may be task-dependent. Finally, accounts that do not call into cause attentional processes cannot be ruled out entirely based on our data. For example, results may be due to different affective reactions in the two groups, with non-smokers implicitly judging NRS stimuli as more aversive, hence the observed pupil dilation. This pattern, not shown in smokers, may be abolished due to increased familiarity with NRS, though its reverse in a constriction would be problematic to explain in these terms. The individual attitude toward addiction-relevant stimuli can be a powerful determinant of the attentional biases reported in literature, and do not exclude the presence of other attentional or perceptual mechanisms. Unfortunately, when compared to other physiological signals, changes in pupil size are relatively slow, which limits the conclusions that can be drawn based on the latency of the effects. We do note, however, that attentional orienting accounts appear partially supported by the correlations with the results of the DPT task, whose parameters are more classically interpreted as spatial attentional biases. Even though there was no significant difference in dwell times for NRS in smokers, the preference for NRS correlated with a more vigorous *constriction* of the pupil in a subsequent, independent task (PV). The opposite was true when assessing the directional bias toward NRS (i.e., the tendency to fixate NRS stimuli first). Pupil constriction was stronger for participants who favored, in the DPT task, first gazing away from NRS stimuli. Previous studies have shown that behavioral performance at short vs. long SOAs can be anticorrelated (della Libera et al., 2019), showing that different mechanisms of attentional engagement can unfold at a different pace. One possibility is that the autonomic signal conveyed by the pupil may reflect the combination of all these underlying processes, i.e. both short-term NRS avoidance and long-term NRS preference. This is in agreement with the idea that pupil size may provide an integrated readout of different attentional networks, perceptual, cognitive, and emotional processes (Strauch et al., *in press*), hence the potential for a superior capability to predict smoking status.

Whilst possibly task-dependent, indeed, change in pupil size to NRS in this setting retained good predictive validity. The information conveyed by pupil size appears necessary and sufficient for a good classification of smoking status (65% accuracy), although additional proxy measures (i.e., RTs gain in the DPT) may improve the classifier’s performance up to 75%. There is substantial room for improvement, both in terms of the choice of the modeling approach and the range of behavioral measures to feed to the classifier. For example, several different SOAs may be needed to exploit the full potential of this paradigm (e.g. Della Libera et al., 2019). However, our results put forward pupil size as a particularly robust and sensitive candidate to this aim. We maintain that the focus on prediction performance instead of classic significance testing represents a leap forward in the attempt to characterize robust biomarkers of addiction as well as cognitive effects (Yarkoni & Westfall, 2017). This also enables one to study the instances in which the classifier does not correctly predict the smoking status of individuals (i.e., false negatives). Interestingly, this happened more often with individuals who smoked more cigarettes per day, i.e. a proxy for the intensity of smoking. In other words, the classifier worked best, and the pupils constricted more vigorously, for participants who smoked less. Previous studies also found increased dwell times for NRS at lower levels of nicotine dependence (Mogg et al., 2003, 2005). It has been proposed that, as dependence increases, the strength of NRS as incentives actually diminishes and leaves space for more habit-driven, automated modes of consumption (Mogg et al., 2005). Our results fit well with these proposals (Di Chiara, 2000), though ours extend to the autonomic response conveyed by pupil size. The major drawback is that, if true, the quest for potential biomarkers or objective indices of dependence should move past indices of attentional bias, as these may be less relevant in heavy(er) smokers. Similarly, this may detract from the idea that treatments aimed at modifying attentional bias can be effective. Besides the nontrivial choice of a paradigm that is capable to bias attention for a sustained period of time, in ecological conditions, and all while maintaining a short administration time, attentional bias may not be the most critical feature of smokers seeking treatment. On the other hand, however, attentional bias may remain useful in characterizing the stage of dependence, potentially triggering different management strategies. There is, furthermore, an alternative reading of these results. Light smokers also refrained from smoking for longer before the experiment, and we did find some evidence for attentional bias being stronger with abstinence time, which may suggest some relationship with increased craving. While we did not observe correlations with the more established tools to measure craving (i.e., the QSU questionnaire), this may be due to the implicit nature of autonomic response vs. the explicit nature of self-report questionnaires dissociating. However, because abstinence time was correlated with the intensity of nicotine dependence – in that people who smoked less were more likely to refrain from smoking for longer times, whereas those who smoked more cigarettes per day were more likely to smoke in the last useful time slot before the experiment – the collinearity in the data prevents strong conclusions in this regard. More research will help shed light on this matter, but we suggest that measuring attentional bias may provide a useful, objective measure for either craving or dependence stage. Pupil size, in particular, appears to be in a more than ideal position to this aim, in that it is capable to jointly reflect affective, perceptual/attentional, and cognitive processes, thus returning an appropriately multifaceted picture.

## 5. Acknowledgments

We are grateful to Dilara Aladağ for help in data collection. E.B. was supported by a grant from the Department of General Psychology, University of Padua, through the program “Dipartimenti di Eccellenza (art.1, commi 314-337 legge 232/2016)” funded by the Italian Ministry of University and Research.

## 6. Declarations

### Funding

This study was carried out within the scope of the research program Dipartimenti di Eccellenza (art.1, commi 314-337 legge 232/2016), which was supported by a grant from MIUR to the Department of General Psychology, University of Padua.

### Conflict of interest

The authors declare no conflict of interest.

### Ethics approval

The study was approved by the Ethics Committee of the University of Padua (protocol n° 3568). Informed consent was obtained from all individual participants included in the study, in accordance with the declaration of Helsinki.

### Open practices statement

The study was not formally preregistered. The data, code, and materials for this manuscript are available in the Open Science Framework repository: https://osf.io/6r5ch/

## Notes

https://osf.io/6r5ch/

### Competing Interest Statement

The authors have declared no competing interest.

### Summary of Updates

Amended following revision in Psychonomic Bulletin & Review. These changes consist in a more precise terminology, which should hopefully get rid of all sources of confusion, an overall linguistic revision, and the inclusion of a much more nuanced discussion of our findings, which should better link them with the relevant theory and better detail one possible mechanistic model for the effects at hand.

